# Associative learning reconfigures colour-guided behaviour in *Drosophila* larvae

**DOI:** 10.64898/2026.10.28.762446

**Authors:** Ali Asgar Bohra, HaDi MaBouDi, Joni Kemppainen, Mikko Juusola

**Affiliations:** School of Biosciences, University of Sheffield, Sheffield S10 2TN, UK; Neuroscience Institute, University of Sheffield, Sheffield S10 2TN, UK; Alfrisin Technologies, 89400 Hyrynsalmi, Finland

## Abstract

Colour-guided behaviour can be innate or learned, yet how a numerically simple nervous system assigns value to spectral cues remains unresolved. We developed an automated differential-conditioning assay in which third-instar *Drosophila* larvae were trained with blue and green light, one of which was paired with electric shock. Wild-type larvae subsequently avoided the shock-paired colour, whereas naïve controls showed no strong blue-green preference. This learned avoidance persisted when the relative intensity cue was transferred to the alternative colour, indicating that behavioural choice followed spectral identity rather than simply tracking the dimmer stimulus. Disruption of either the blue-sensitive Rh5 or green-sensitive Rh6 photoreceptor pathway significantly weakened the training-induced behavioural shift relative to wild type, with Rh5 mutants showing only a small residual shift. Complementary extracellular recordings confirmed light-evoked activity in the larval visual system, with robust responses to ultraviolet and broadband white illumination but smaller and more variable responses to blue and green light. Together, these results show that larvae can discriminate blue from green, demonstrating elementary colour vision; that associative experience reconfigures their colour-guided behaviour; and that both photoreceptor pathways contribute to this learned response. This paradigm separates learned spectral preference from baseline light preference and provides a tractable framework for linking defined sensory inputs to the circuit mechanisms of visual memory.

## Introduction

Learning allows sensory cues to acquire predictive value and guide behaviour beyond innate preferences. Colour provides a stringent test of this capacity because photoreceptor excitation depends jointly on spectral composition and intensity; unless these dimensions are dissociated, apparent colour discrimination may instead reflect differences in stimulus intensity rather than spectral identity^1,2^. In adult *Drosophila* and bees, behavioural studies have demonstrated learned colour discrimination by combining differential conditioning with intensity controls, while physiological and genetic studies have revealed how comparisons between spectrally distinct photoreceptor channels contribute to chromatic processing^3-7^. These findings establish that learned colour-guided behaviour depends not only on spectral sensitivity and the comparison of photoreceptor signals but also on associative processes that assign behavioural value to chromatic information^1,8^. Whether the numerically compact larval *Drosophila* visual system can discriminate spectral cues through associative learning, rather than relying exclusively on innate wavelength preferences or differences in stimulus intensity, however, remains unresolved.

*Drosophila* offers extensive genetic and circuit-level access to this problem. Adult flies discriminate spectral stimuli through the combined activity of multiple photoreceptor classes and interacting chromatic and achromatic pathways^6,7^. Circuit-level studies have further demonstrated that recurrent interactions within the adult optic lobe contribute to the transformation of photoreceptor signals into hue-selective neuronal responses^1^. Complementing these circuit-level findings, aversive-conditioning experiments suggest that distances within a multichannel, opponent photoreceptor space can partly predict discrimination performance across wavelengths^7^. However, adult spectral processing is distributed across the complex visual system, making it challenging to trace the transformation of spectral signals from sensory encoding to associative memory and behavioural choice. The larval visual system offers a simpler preparation, in which each Blowing organ contains only 12 photoreceptors belonging to two opsin-defined classes ^9^. Moreover, a synaptic-resolution reconstruction of the first-instar larval brain, comprising 3,016 neurons, provides an anatomical framework for tracing sensory pathways towards learning and behavioural circuits^10^. This organisation provides an opportunity to investigate how spectral information is processed and assigned learned value within a compact, experimentally and genetically accessible nervous system.

In each larval eye, or Bolwig organ, four photoreceptors express blue-sensitive Rhodopsin 5 (Rh5), whereas eight express green-sensitive Rhodopsin 6 (Rh6)^9,11^. Their axons terminate in the larval optic neuropil, where synaptic reconstruction reveals distinct but partially convergent pathways to downstream visual interneurons^9,11^. Importantly, connectomic reconstructions have identified anatomically defined pathways linking the larval optic neuropil to the mushroom body. Three visual projection neurons, the fifth lateral neuron (5th-LaN) and two non-clock lateral neurons (nc-LaNs), form synaptic connections with Kenyon cells in the mushroom-body accessory calyx^12,13^. This organisation provides a potential route through which visual information could reach associative-learning circuits. However, whether these pathways support spectral discrimination and the assignment of learned value to colour cues remains unknown. Rh5 photoreceptors dominate rapid light avoidance under standard laboratory conditions, whereas either photoreceptor subtype can support circadian entrainment^14^. Larval phototaxis involves active temporal sampling of light during crawling and head casts^15^and experiments under natural illumination reveal a context-dependent contribution of Rh6 to light avoidance^16^. More specifically, Rh5- and Rh6-associated pathways contribute differently to spatial and temporal processing of visual cues, respectively^17^. Earlier behavioural studies demonstrated wavelength-dependent light responses in wild-type and mutant *Drosophila* larvae, establishing spectral sensitivity without determining whether larvae can acquire learned preferences between distinct colours^16,18^. These functional differences raise the question of how the two photoreceptor populations contribute to the physiological encoding of spectral stimuli and whether their distinct responses support learned colour-guided behaviour.

Previous work established that *Drosophila* larvae can associate light or darkness with appetitive and aversive outcomes^19,20^ and that electric shock can serve as an effective unconditioned stimulus for larval associative learning^21,22^. However, learning to distinguish light from darkness does not establish discrimination between spectral cues, while wavelength-dependent phototaxis alone cannot reveal whether colour acquires learned value through experience. Addressing this distinction requires conditioning protocols that reciprocally pair blue and green stimuli with reinforcement, account for pre-existing preferences, and determine whether learned behaviour follows spectral identity when relative stimulus intensities are reversed. Manipulating Rh5 and Rh6 can then test whether these pathways contribute differently to learned spectral discrimination and innate light avoidance. Combined with synapse-resolution reconstructions of the larval visual system, mushroom body, and first-instar brain^11,23,24^, this approach provides a foundation for future investigations of how spectral information becomes associated with reinforcement and guides behavioural choice.

Here, we developed an automated differential-conditioning assay in which blue and green illumination alternated, with one colour paired with electric shock. Wild-type larvae subsequently avoided the shock-associated colour, and this learned shift persisted when the relative-intensity cue was reversed at test. Disruption of either Rh5 or Rh6 significantly attenuated the training-induced behavioural shift relative to wild type, despite near-neutral innate blue–green choice in all three genotypes. Complementary electroretinogram (ERG) recordings confirmed light-evoked responses in wild-type and Rh5- and Rh6-mutant larvae, revealing differences in their responses under the tested illumination conditions. Together, these findings demonstrate learned blue–green discrimination that cannot be explained solely by the tested relative-intensity cue and implicate both photoreceptor pathways in normal colour-guided learning. This combination of behavioural, genetic and physiological evidence establishes a foundation for investigating how spectral information is processed and acquires learned behavioural value within the compact nervous system of *Drosophila* larvae.

## Results

### An automated assay measures differential aversive colour learning

We asked whether *Drosophila* larvae, whose Bolwig organs contain only two opsin-defined photoreceptor populations expressing Rh5 and Rh6, can discriminate blue from green and learn to avoid a colour associated with electric shock. To test this, we developed a programmable apparatus that presented blue and green illumination from below a 90-mm agarose arena and synchronised electric shock with the reinforced colour (**Figure 1a**). We first measured baseline colour preference by testing naïve late third-instar larvae for 60 s in a half-blue, half-green arena. Independent groups then underwent five training cycles, each comprising 30 s of blue and 30 s of green illumination. One colour served as the reinforced conditioned stimulus (CS+) and was paired with shock, whereas the other served as the unreinforced conditioned stimulus (CS−), with reciprocal colour assignments across groups. Immediately after training, larvae were transferred to a fresh two-choice arena and tested for 60 s without reinforcement (**Figure 1b**, see method).

**Figure 1.**
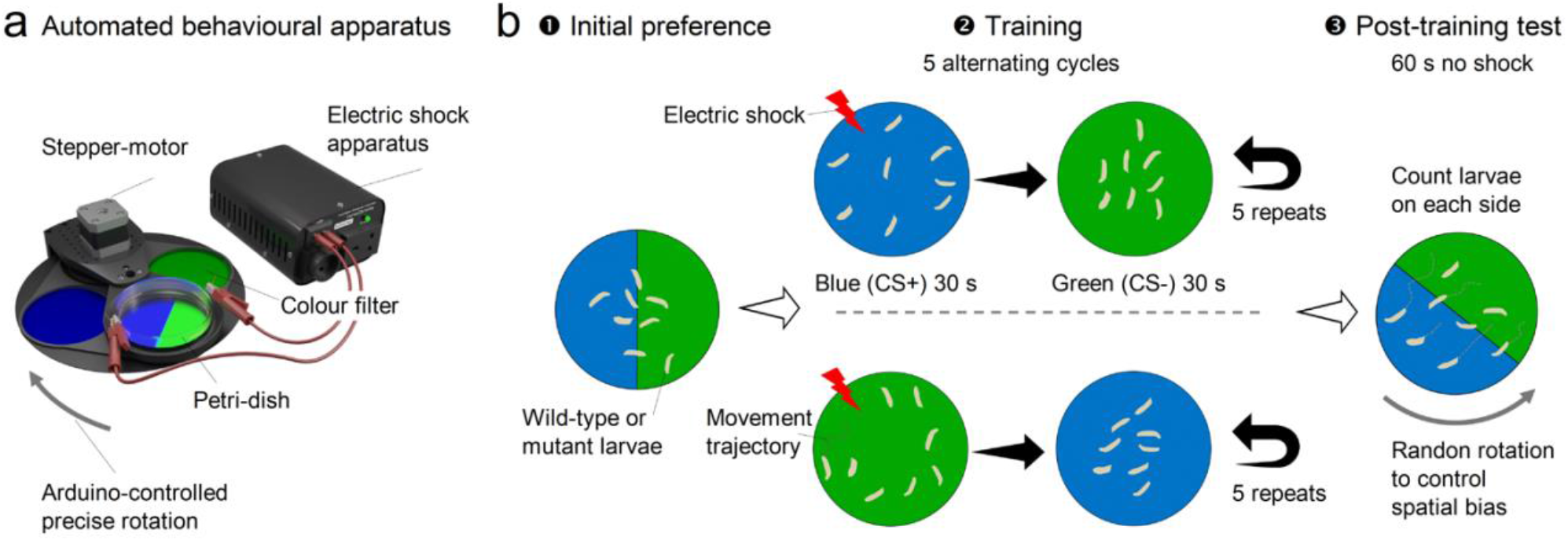
Automated apparatus and experimental protocol for aversive colour learning in *Drosophila* larvae. (**a**) Custom-built automated behavioural apparatus. A Petri dish containing 1% agarose was positioned above interchangeable blue and green optical filters and was illuminated from below. An Arduino-controlled stepper motor enabled precise rotation of the filter platform, while a relay system synchronised colour presentation with electrical-shock delivery. (**b**) Experimental sequence. (1) Initial preference: naïve third-instar larvae were placed at the centre of an arena divided equally into blue- and green-illuminated halves and allowed to move freely for 60 s without reinforcement. Larvae on each side were then counted. (2) Training: larvae underwent five alternating cycles comprising a 30-s presentation of the conditioned stimulus (CS+) paired with electrical shock and a 30-s presentation of the unreinforced stimulus (CS−). Blue and green were assigned reciprocally as the CS+ across experimental groups. (3) Post-training test: immediately after training, larvae were transferred to a fresh arena divided into equal blue and green halves and tested for 60 s without reinforcement. The number of larvae on each side was counted, and the arena orientation was randomised between test trials to control for spatial bias. Larval positions and movement trajectories are shown schematically.

Baseline and post-training measurements used independent groups of larvae, rather than paired measurements of the same groups. The primary standard-intensity dataset comprised 104 groups and 3,971 responding larvae: 40 groups of wild type and 32 groups of each mutant genotype (Rh5^2^ and Rh6^1^). Because larvae within one dish shared experimental conditions, including handling, visual stimulation and reinforcement, the larval group was treated as an independent biological replicate. Statistical inference was therefore based on group-level choice count analysed using beta-binomial mixed-effects models, with experimental batch included as a random intercept (see Method for details).

### Wild-type larvae acquire robust avoidance of the shock-associated colour

Wild-type (WT) larvae showed a marked training-induced shift in preference away from the colour paired with electronic shock (CS+; **Figure 2a**). The mean conditioned-colour avoidance (negative preference index) decreased from 0.052 ± 0.029 in naïve control groups to -0.216 ± 0.034 after conditioning, indicating that trained larvae were less likely to choose the shock-associated colour.

**Figure 2.**
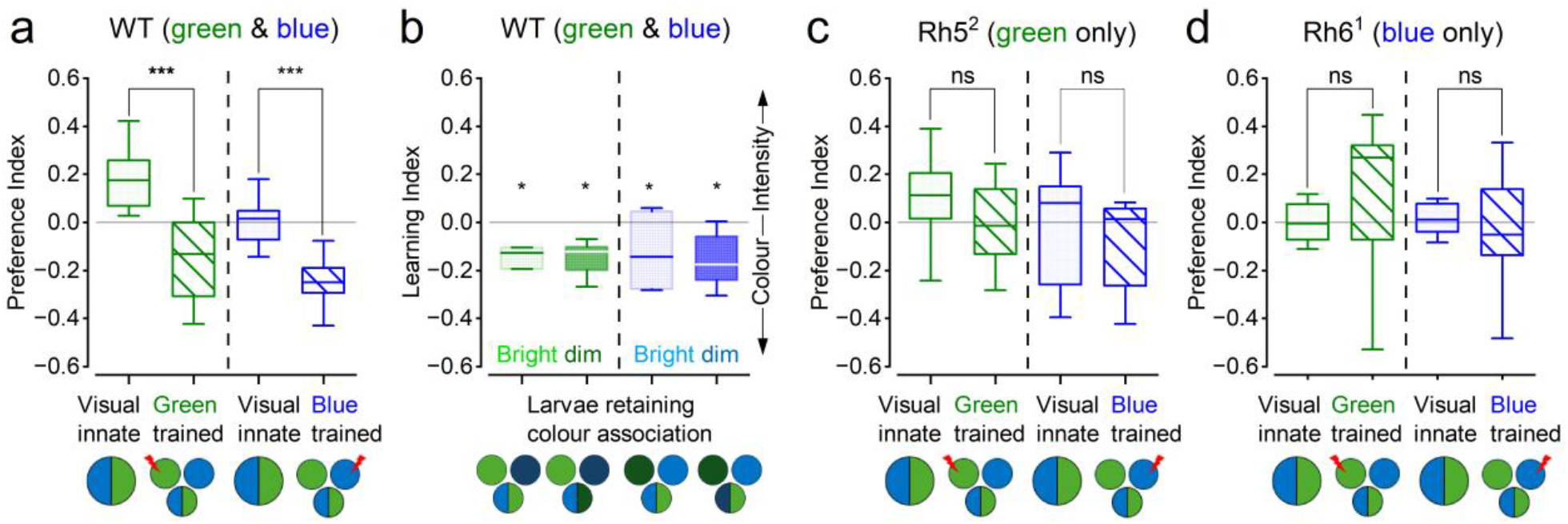
Learned colour avoidance in wild-type larvae and its attenuation in Rh5 and Rh6 mutants. (**a**) Wild-type (WT) larvae, possessing blue-sensitive Rh5 and green-sensitive Rh6 photoreceptors, shift their preference away from the colour paired with electric shock (CS+), whether green or blue. (**b**) WT larvae distinguish green from blue and retain the learned colour association across the tested intensity changes. After training with one colour at about one-third of its original intensity, avoidance (negative preference) persists when both colours are restored to their original intensities or when the reduced-intensity cue is transferred to the alternative colour. Thus, learned avoidance is colour-specific and cannot be explained solely by avoidance of the dimmer stimulus. (**c, d**) Rh5^2^ mutants (green-sensitive channel only; **c**) and Rh6^1^ mutants (blue-sensitive channel only; **d**) show weaker training effects than WT larvae and neither within-genotype effect reaches significance. These results support attenuated colour-associated learning rather than an absolute inability to learn. Independently analysed innate blue-green choice is close to neutral in all three genotypes.

We then asked whether the magnitude of the training effect varied depending on whether blue or green served as the conditioned stimulus (CS+). A full training × genotype × conditioned-colour model did not improve fit over the more parsimonious training × genotype + conditioned-colour model (likelihood-ratio χ^2^ = 3.246, df = 5, p = 0.662). We therefore estimated planned training-effect contrasts from the parsimonious model, accounting for differences in conditioned-colour choice associated with CS+ identity.

The beta-binomial mixed model showed that training reduced the odds of choosing the conditioned colour by approximately 41% in wild-type larvae relative to naïve controls (odds ratio [OR] = 0.586, 95% confidence interval [CI] 0.465–0.740, Holm-adjusted p = 2.05 × 10^−^5; **Figure 2a**). Together with reciprocal CS+ assignments, this training-dependent reduction in conditioned-colour choice supports learned avoidance of the shock-associated stimulus, although non-associative effects of shock exposure cannot be fully excluded. We next tested whether this avoidance was guided by spectral identity of differences in stimulus intensity.

### Learned avoidance follows spectral identity when the relative-intensity cue is reversed

To determine whether conditioned avoidance followed spectral identity rather than relative stimulus intensity, we performed a separate wild-type (WT) experiment in which the relationship between colour and intensity was altered between training and testing. During asymmetric training, either blue or green was presented at approximately one-third of its original intensity, with conditioned-colour assignments counterbalanced across groups. At test, both colours were either restored to their original intensities, or the reduced-intensity cue was transferred to the alternative colour (**Figure 2b**).

Conditioned-colour avoidance persisted under both test conditions. The estimated probability of choosing the conditioned colour was below 0.5 when both colours were restored to their original intensities (p[CS] = 0.432, 95% CI 0.398–0.468, Holm-adjusted p = 0.00042) and remained below 0.5 when the reduced-intensity cue was transferred to the alternative colour (p[CS] = 0.436, 95% CI 0.401–0.471, Holm-adjusted p = 0.00042; **Figure 2b**). Cue reversal had no detectable effect on conditioned-colour choice (OR = 1.014, 95% CI 0.829–1.239, p = 0.896). A full model including interactions between test mode, conditioned colour and reduced-intensity assignment did not improve fit over an additive model (likelihood-ratio χ^2^ = 1.481, df = 4, p = 0.830).

Green-conditioned groups showed a modestly higher probability of choosing the conditioned colour than blue-conditioned groups (OR = 1.283, 95% CI 1.049–1.569, Holm-adjusted p = 0.031), whereas the colour assigned reduced intensity during training had no detectable effect (OR = 1.048, 95% CI 0.855–1.285, adjusted p = 0.652). Together, these findings show that learned avoidance remained directed towards the shock-associated colour despite changes in relative stimulus intensity. Thus, conditioned choice cannot be explained solely by avoidance of the lower-intensity stimulus under the conditions tested.

### Rh5 and Rh6 mutants show attenuated learned avoidance

Does colour-associated learning depend on input from both photoreceptor types? To address this question, we tested larvae lacking functional Rh5 or Rh6, retaining Rh6- or Rh5-mediated photoreception, respectively. Each mutant genotype underwent conditioning with either green or blue paired with shock (**Figure 2c, d**).

Rh5 mutants showed a smaller shift in conditioned-colour preference than wild-type (WT) larvae: the mean preference index for the conditioned colour, with negative values indicating avoidance, shifted from 0.041 ± 0.057 in naïve controls to -0.048 ± 0.048 after conditioning. The corresponding training effect was not statistically significant (OR = 0.878, 95% CI 0.671–1.149, Holm-adjusted p = 0.688). Rh6 mutants showed no comparable shift towards conditioned-colour avoidance: mean preference indices were 0.019 ± 0.025 in naïve controls and 0.045 ± 0.073 after conditioning, with no detectable training effect (OR = 1.008, 95% CI 0.776–1.310, adjusted p = 0.950).

To determine whether disruption of either photoreceptor pathway attenuated learned avoidance, we compared the training effects between each mutant and WT larvae. Both genotype-by-training interactions were statistically significant. Relative to the WT training effect, the Rh5 interaction had an OR of 1.497 (95% CI 1.049–2.137, Holm-adjusted p = 0.026), and the Rh6 interaction had an OR of 1.719 (95% CI 1.211–2.440, adjusted p = 0.0048). Interaction odds ratios greater than one indicate that conditioning reduced the probability of choosing the shock-associated colour less strongly in the mutants than in WT larvae. These comparisons support attenuated colour-associated learning in both mutants. However, these results do not establish a complete loss of learning, as the confidence intervals remain compatible with residual training effects.

Across genotypes and training conditions, groups were modestly more likely to choose the conditioned colour when green rather than blue served as CS+ (OR = 1.194, 95% CI 1.029–1.386, p = 0.019). This colour-dependent difference was accounted for in the statistical model and did not account for the genotype-by-training interactions.

### Baseline blue-green preferences are close to neutral

We next examined whether differences in baseline blue–green preference could contribute to the reduced training effects observed in the mutants. To assess this possibility, we analysed the probability that naïve larvae chose blue, irrespective of which colour was designated CS+ in the corresponding conditioning experiment. Model-estimated probabilities of choosing blue were 0.488 in WT larvae (95% CI 0.449–0.527), 0.460 in Rh5 mutants (0.411–0.511), and 0.508 in Rh6 mutants (0.460–0.555). None differed significantly from equal choice (0.5) after Holm correction (all adjusted p ≥ 0.368), and neither mutant differed detectably from WT larvae (both adjusted p = 0.797).

Although these near-neutral baseline preferences do not establish equivalent visual perception across genotypes, they provide no evidence for strong pre-existing blue–green choice biases that could account for the attenuated training effects in the mutants.

### Colour-evoked extracellular responses reveal distinct temporal contributions from the two larval photoreceptor pathways

We adapted the established extracellular electroretinogram (ERG) approach to record light-evoked responses from intact *Drosophila* larvae. The larval visual system differs markedly from that of adult flies. Its two small Bolwig organs contain only 24 photoreceptors in total: eight expressing blue-sensitive Rh5 and 16 expressing green-sensitive Rh6. These photoreceptors feed into a downstream network with multiple neuronal types and transmitter systems (**Figure 3a**). Unlike adult compound eyes, the larval visual organs lack screening pigments, and the surrounding tissues are translucent, allowing illumination to reach the photoreceptors from multiple directions. Larval Rh5 and Rh6 photoreceptors have broad, substantially overlapping spectral sensitivities, whereas adult R8 photoreceptors expressing these same rhodopsins have narrower spectral sensitivities filtered by intraommatidial pigments^25,26^. Consequently, individual colour stimuli can activate both larval receptor types to different degrees (**Figure 3b**).

**Figure 3.**
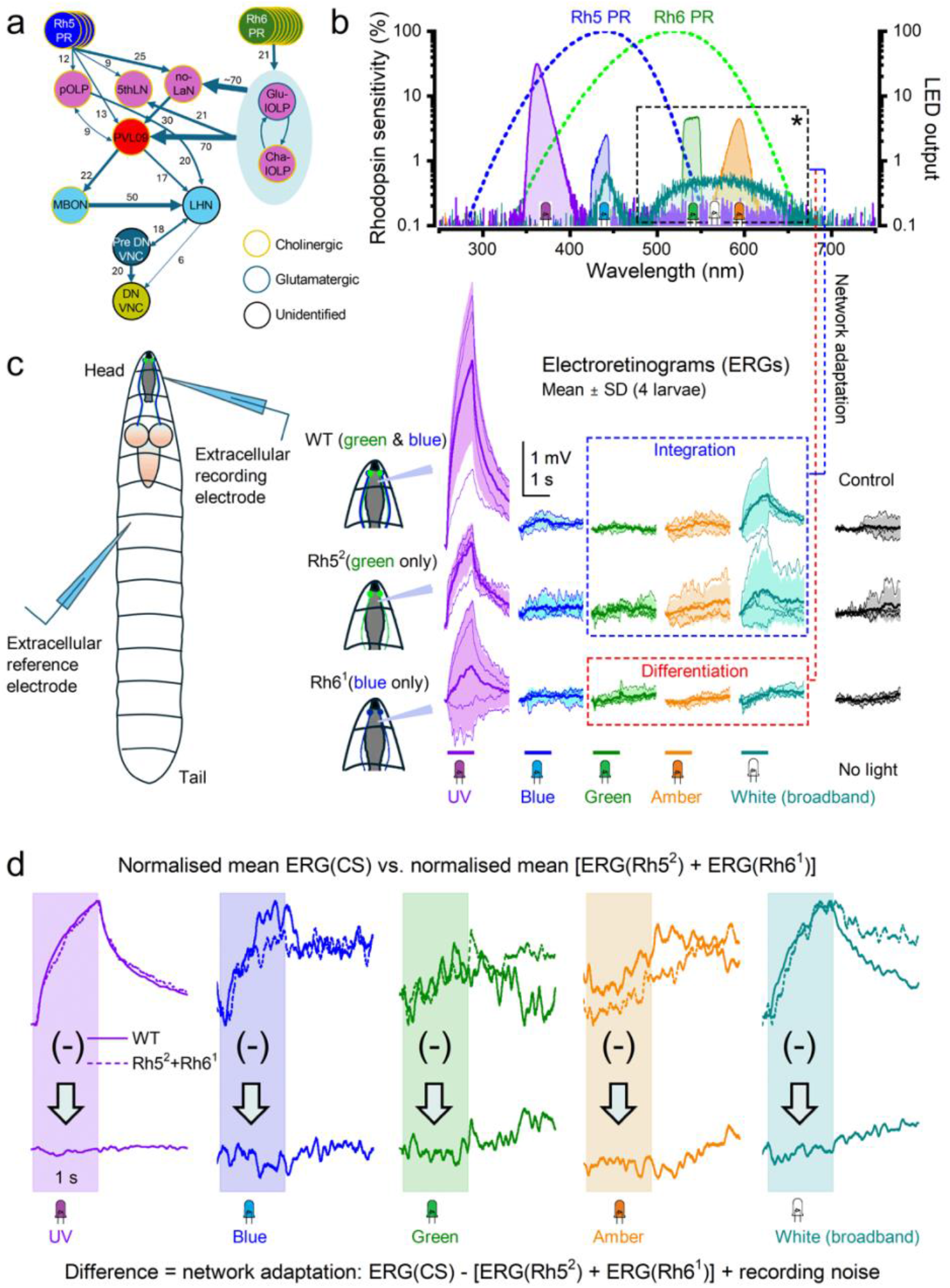
Larval visual circuits generate distinct spectral responses from UV to amber despite expressing only blue- and green-sensitive rhodopsins. **(a)** Wiring diagram of the early larval visual network, showing inputs from four Rh5-expressing (blue-sensitive) and eight Rh6-expressing (green-sensitive) photoreceptors per Bolwig organ. Numbers indicate synapse counts; neuronal outline colours indicate neurotransmitter identity. **(b)** Predicted spectral sensitivities of Rh5 and Rh6 (dashed nomograms; left axis) and measured emission spectra of the UV, blue, green, amber and broadband white LEDs (right axis). **(c)** Left: positions of the extracellular recording electrode near the larval head and the reference electrode towards the posterior body. Right: ERGs evoked by 1-s light pulses in wild-type Canton-S (CS), Rh5^2^ and Rh6^1^ larvae. The mutants retain functional Rh6 (green-sensitive) or Rh5 (blue-sensitive) photopigment, respectively. Traces and shading show mean ± SD (four larvae per genotype); coloured bars indicate stimulus duration. No-light controls are shown on the right. UV stimulation evoked the largest responses in all three genotypes, while the different LEDs elicited distinct response waveforms. Dashed boxes highlight response features interpreted as integration (sustained activity) and differentiation (more transient activity). (d) Comparison of normalised mean wild-type ERGs (solid lines) with the normalised sum of the mean Rh5^2^ and Rh6^1^ mutant ERGs (dashed lines). Lower traces show the difference between these waveforms; shaded regions indicate the 1-s stimuli. Summed mutant responses reproduce most features of the wild-type responses with residual differences likely reflecting phenotype-specific homeostatic interactions and adaptation within the intact visual network, together with recording noise.

We immobilised dark-adapted third-instar larvae with beeswax and positioned a glass-capillary recording electrode against the anterior body surface near the Bolwig organs, with a similar reference electrode against the middle of the body (**Figure 3c**, left). We then recorded extracellular voltage responses to 1-s LED pulses of ultraviolet, blue, green, amber and broadband white light. The recordings revealed light-evoked responses whose amplitudes and time courses varied with stimulus and genotype (**Figure 3c**, right). These extracellular signals may contain contributions from both photoreceptors and downstream neurons; the present recordings do not isolate their relative contributions.

In WT larvae, ultraviolet stimulation produced the largest responses, while broadband white light also evoked a pronounced response relative to the no-light controls. Blue, green and amber stimuli evoked much smaller mean responses. Response amplitudes must be interpreted in relation to both the overlapping photoreceptor sensitivities and the different spectral outputs of the LEDs (**Figure 3b**). The recordings in **Figure 3c** therefore compare responses under the tested illumination conditions rather than providing an intensity-controlled measure of spectral sensitivity.

The WT responses included sustained components that developed during illumination and subsided after light offset. These were particularly apparent under ultraviolet and white illumination; the smaller responses to the other stimuli were more variable. The sustained waveforms are compatible with temporal integration within the visual system (**Figure 3c**, blue dashed box). However, extracellular polarity alone does not establish whether excitation or inhibition dominates the underlying cellular activity.

Both Rh5^2^ and Rh6^1^ mutants retained light-evoked responses, but their waveforms differed. Rh5^2^ mutants lack functional Rh5 and retain “green-sensitive” Rh6-mediated photoreception, whereas Rh6^1^ mutants lack functional Rh6 and retain “blue-sensitive” Rh5-mediated photoreception. Thus, across the two Bolwig organs, Rh5^2^ mutants retain Rh6-mediated light detection in 16 photoreceptors, whereas Rh6^1^ mutants retain Rh5-mediated light detection in eight.

Rh5^2^ mutant responses generally resembled the WT waveforms, particularly under ultraviolet and white illumination, but their mean amplitudes were smaller. Their sustained components are consistent with a contribution from the Rh6 pathway to the more slowly varying responses observed in WT larvae (**Figure 3c**, blue dashed box). However, differences in extracellular amplitude cannot be attributed solely to the number of functional photoreceptors.

Rh6^1^ mutants showed smaller mean responses and different temporal profiles. In particular, the green, amber and white responses included brief deflections around stimulus transitions that contrasted with the more sustained responses of Rh5^2^ mutants (**Figure 3c**, red dashed box). These differences suggest that the Rh5 and Rh6 pathways make distinct temporal contributions to the extracellular response, potentially combining more transient and more sustained components. An inhibitory or differentiating contribution from the Rh5 pathway is one possible interpretation, but these recordings alone cannot establish it.

Finally, we asked whether adding the corresponding mean responses of the two mutants could approximate the WT waveform for each stimulus (**Figure 3d**). Because extracellular response amplitudes can vary with electrode placement and recording conditions, we compared the amplitude-normalised WT mean with the amplitude-normalised sum of the two mutant means. This comparison assesses waveform similarity rather than additivity of absolute response amplitudes.

The summed mutant responses approximated the WT waveform particularly closely for ultraviolet stimulation and captured some temporal features of the responses to the other stimuli. Nevertheless, differences remained, including during recovery after light offset. The lower traces in **Figure 3d** show these residual differences, which could reflect interactions between the two pathways in WT larvae, genotype-dependent homeostatic adaptation or compensation, and variability associated with recording and normalisation. Thus, summing the mutant responses partially reconstructs WT response dynamics. This resembles findings in adult flies, where summing ERG responses from the R1–R6 and R7/R8 photoreceptor populations broadly reproduces the WT ERG^26^. In larvae, however, mutation-specific network adaptation may also shape the recorded responses.

## Discussion

Our results provide evidence for learned spectral discrimination in *Drosophila* larvae, demonstrating that associative experience can change the behavioural meaning of spectral information in this numerically simple nervous system. This capacity is influenced by the integrity of two broadband photoreceptor types with strongly overlapping spectral sensitivities. WT larvae learned to avoid either blue or green when that colour predicted electric shock. This learned avoidance cannot readily be explained by innate wavelength preference or the tested relative-intensity cue alone: the conditioned colour was assigned reciprocally, naïve larvae showed no strong blue–green preference, and avoidance remained directed towards the conditioned colour when the relative-intensity cue was transferred to the alternative stimulus. Together, these findings support experience-dependent assignment of aversive value to spectral cues, beyond the preferences observed in naïve larvae.

The intensity-transfer experiment provides the strongest evidence for learned discrimination of spectral cues beyond the tested relative-intensity relationship. When colour identity and relative intensity predicted opposing choices, larvae continued to avoid the colour paired with shock. This argues against a strategy based exclusively on avoiding the lower-intensity stimulus under the conditions tested. Nevertheless, demonstrating more general intensity-invariant colour discrimination would require behavioural tests across independently varied stimulus intensities, together with estimates of photoreceptor quantum catches and potential achromatic cues^2,27^. Spatial non-uniformity, adaptation and unequal photoreceptor excitation could still provide residual achromatic information^27^. Spatial non-uniformity, adaptation and unequal receptor quantum catch could still provide residual achromatic information. Our results therefore demonstrate a learned spectral association that survives reversal of the tested relative-intensity cue, while leaving the extent of intensity invariance unresolved. Comparable reciprocal conditioning and intensity controls have been central to demonstrations of colour discrimination in adult flies and bees^2,4,6,28^.

### From photoreceptors to learned colour

The organisation of the insect visual pathway suggests where this spectral comparison might arise. In adult *Drosophila*, broadband R1-R6 photoreceptors project to the lamina, whereas spectrally specialised R7 and R8 photoreceptors project principally to the medulla. These pathways are not independent: inner photoreceptor signals can influence nominally achromatic pathways, and interactions between chromatic and achromatic circuits contribute to colour and motion processing^6,26,28^. Recent studies further demonstrate that recurrent interactions within the adult optic lobe contribute to the emergence of hue-selective responses, illustrating how circuit computations can transform broadly tuned sensory inputs into more selective chromatic representations^1^.

The larval pathway is anatomically simpler and should not be described using the adult lamina– medulla sequence. Rh5- and Rh6-expressing photoreceptors of the Bolwig organ project directly through the Bolwig nerve to the larval optic neuropil. Connectomic reconstruction reveals parallel but partially convergent visual pathways within and beyond this neuropil^8,9^. Rh5 dominates rapid light avoidance under standard laboratory conditions, whereas Rh6 contributes more strongly under particular natural illumination conditions and to visual computations based on temporal changes during larval movement and head casting^14-16,29^.

The present mutant results suggest that pathways previously implicated in innate visual navigation also contribute to learned colour-guided behaviour. Disruption of either Rh5 or Rh6 attenuated the wild-type training effect, but the present experiments do not establish whether this attenuation reflects impaired spectral discrimination, altered visual sensitivity or other effects of the mutations. Instead, the two pathways may jointly generate a comparative spectral code, or may make complementary contributions to adaptation, gain regulation and temporal sampling. Distinguishing these possibilities will require measurements of the visual responses and behavioural discriminability supported by each pathway^26^.

Because Rh5 and Rh6 have broad, overlapping sensitivity functions, loss of either receptor alters the balance and reliability of the remaining population response rather than removing an isolated detector for one colour^30^. Near-neutral innate choice in both mutants provides no evidence for a strong baseline preference that would explain the attenuated behavioural effect, but does not demonstrate normal sensory encoding. Cell-specific rescue, acute silencing and cell-type-specific physiological measurements will be required to distinguish reduced spectral discriminability from developmental compensation or genetic-background effects^31^.

### Distinct temporal contributions to larval visual processing

The extracellular recordings complement the behavioural results by showing that both mutant pathways retain light responsiveness but differ in response amplitude and temporal profile. Rh6-mediated responses in Rh5^2^ mutants generally resembled the more sustained WT waveforms, whereas Rh5-mediated responses in Rh6^1^ mutants included more transient components around stimulus transitions. These differences suggest that the two pathways contribute complementary temporal information, potentially supporting the integration and detection of changes in visual input during larval movement and head casting.

Summing the normalised mutant responses partially reconstructed WT response dynamics, particularly under ultraviolet stimulation. The remaining differences are compatible with pathway interactions and mutation-specific adaptation or compensation, alongside recording variability. Thus, the behavioural deficits occur despite retained light responsiveness and may reflect altered spectral comparison, temporal processing or both. However, the extracellular recordings neither isolate the contributing cells nor establish chromatic opponency, and the unequal LED outputs prevent intensity-controlled comparisons of spectral sensitivity. Together, the behavioural and physiological results support complementary contributions from Rh5 and Rh6 pathways to learned spectral choice, while leaving the underlying circuit computations unresolved.

### A candidate circuit for colour memory

A candidate circuit for learned colour discrimination can be proposed from the established organisation of the larval visual system. Rh5- and Rh6-dependent signals are processed through distinct but partially convergent pathways within the larval optic neuropil, where local interneurons may transform their activity before visual information is relayed to higher brain centres^12^. Three identified visual projection neurons, the fifth lateral neuron (5th-LaN) and two non-clock lateral neurons (nc-LaNs), project from the larval optic neuropil to the mushroom-body accessory calyx, where they form synaptic connections with Kenyon cells^12,13^. These pathways could therefore convey spectral information to associative circuits, where coincidence with reinforcement signals may alter the behavioural value of visual stimuli^22^. Whether these neurons encode discriminable colour information or are required for the colour-associated learning observed here remains to be established.

In adult *Drosophila*, visual and olfactory associative memories can recruit shared mushroom-body compartments and dopaminergic reinforcement pathways^32^. In adult *Drosophila*, anatomically distinct visual projection pathways convey chromatic and brightness information to mushroom-body Kenyon cells and make different contributions to learned colour and intensity discrimination^33^. More recent connectomic analyses have further revealed a specialised subset of Kenyon cells receiving predominantly visual inputs^34^. A parsimonious hypothesis is therefore that colour-responsive projection neurons activate a sparse Kenyon-cell population, while aversive reinforcement-related dopaminergic pathways provide signals capable of modifying colour-associated synaptic representations^35,36^. Plasticity at Kenyon-cell-to-mushroom-body-output-neuron synapses could then alter the behavioural value assigned to the conditioned colour.

Evidence from bees supports the general architecture of this hypothesis. Bee photoreceptor signals are progressively transformed through the lamina, medulla and lobula, where colour-opponent and higher-order visual responses emerge. Projection neurons from the optic lobes innervate the visual collar of the mushroom-body calyx, and many of these mushroom-body input neurons are colour-sensitive^4,37^. Individual performance in bumblebee colour discrimination has also been associated with microglomerular organisation in the mushroom-body collar^38^. These findings suggest a conserved organisational principle in which optic-lobe circuits process spectral information before delivering it to associative networks that assign learned value.

The comparison should nevertheless remain cautious. Visual learning in insects is not uniformly mushroom-body dependent. In adult flies, the required circuit depends on the task: some visual associative memories recruit the mushroom bodies, whereas visual place learning depends on central-complex circuitry, and several earlier visual-learning paradigms were retained after mushroom-body disruption^32,33,39,40^. The circuit underlying larval blue–green conditioning must therefore be determined experimentally rather than inferred from adult flies or bees.

Where is the larval colour association stored? The present experiments do not test this directly. The larval optic neuropil is now mapped at synaptic resolution^11^ and the complete larval mushroom-body and whole-brain connectomes provide defined routes for tracing sensory convergence, dopaminergic reinforcement, and output pathways^23,41^. In adults, visual and olfactory memories use partly shared mushroom-body circuitry^32^. Although anatomical pathways linking the larval visual system to mushroom-body Kenyon cells have been established ^12,13^, their functional contribution to colour-associated learning remains unresolved. In particular, it is unknown whether these visual projection neurons encode differences between blue and green stimuli, whether associative plasticity modifies their downstream representations, or whether mushroom-body circuits are necessary for the learned avoidance observed here.

Targeted experiments can now test this pathway directly. Silencing candidate larval visual projection neurons during acquisition or retrieval would establish whether they carry the conditioned stimulus. Manipulating projection neurons and/or local optic interneurons, aversive dopaminergic neurons, and mushroom-body output neurons would then test associative convergence and memory expression. Calcium imaging or electrophysiology across Rh5 and Rh6 photoreceptors, larval optic-neuropil interneurons and higher-order targets could reveal where blue and green responses first become separable and where conditioning changes stimulus value. The larval visual, mushroom-body and whole-brain connectomes provide an anatomical framework for prioritising these experiments^11,23,24^.

The immediate post-training test demonstrates acquisition and short-term expression, but not long-term memory. Earlier studies established that larvae could associate light or darkness with reinforcement and that electric shock is an effective aversive unconditioned stimulus^19,21^. The present paradigm advances this work by requiring a choice between spectral cues and by dissociating colour from the tested intensity relationship. Retention, reversal and extinction experiments will be needed to establish the persistence and flexibility of the memory. Tracking individual trajectories would additionally reveal whether learning changes run duration, turning probability, head-cast direction or boundary-crossing decisions.

In summary, associative experience reconfigures larval colour-guided behaviour. Wild-type larvae learn to avoid either blue or green when it predicts punishment; this response follows colour after reversal of the relative-intensity cue, and both Rh5 and Rh6 pathways contribute to the full learning effect. The results distinguish learned spectral choice from the Rh5-dominated organisation of innate phototaxis and establish a tractable route from two defined photoreceptor populations, through a compact first-order visual neuropil, to candidate reinforcement and memory circuits. This combination of controlled behaviour, genetics and synapse-resolution anatomy makes larval colour learning a powerful system for determining how a small brain converts spectral input into learned action.

## Limitations of the Study

The behavioural assay measured group-level colour preference immediately after conditioning and therefore does not resolve individual learning trajectories or distinguish acquisition from short-term memory expression. Although reciprocal colour–shock conditioning and intensity-transfer tests support learned spectral discrimination, the absence of explicitly unpaired, shock-only and colour-only controls prevents complete exclusion of non-associative effects of stimulus exposure or sensitisation. Furthermore, the intensity-transfer experiment demonstrates that conditioned avoidance persists after reversal of the tested relative-intensity cue but does not establish complete intensity-invariant colour vision. Residual achromatic information arising from differences in photoreceptor quantum catch, adaptation or spatial illumination cannot be excluded. Future experiments incorporating independently varied photon fluxes, receptor-based spectral modelling and additional reinforcement controls will be necessary to establish the generality and specificity of learned colour discrimination.

Disruption of either Rh5 or Rh6 attenuated conditioned-colour avoidance relative to wild type, but these effects cannot be attributed specifically to deficits in associative learning. Changes in spectral sensitivity, sensory discriminability, visual adaptation or developmental organisation may also contribute. Similarly, near-neutral baseline preferences do not establish equivalent sensory encoding across genotypes. Genetic-background differences and partially distinct experimental periods may further influence comparisons^31^. Cell-specific rescue, acute photoreceptor manipulation and independent behavioural measures of spectral discrimination will be required to determine the causal contributions of each pathway to sensory encoding and learned choice.

Finally, the ERG recordings provide complementary evidence of light-evoked physiological responsiveness in wild-type and photoreceptor-mutant larvae but remain exploratory. The limited number of preparations, variability in electrode placement and unequal spectral outputs of the light sources constrain quantitative comparisons across genotypes and stimuli. Moreover, the ERG signals do not isolate photoreceptor activity from downstream neural contributions, nor do they establish specific temporal computations, chromatic opponency or the physiological mechanisms underlying colour-associated learning. Accordingly, the recordings should be interpreted as preliminary characterisation of larval visual responsiveness rather than direct evidence for pathway-specific computations or associative memory mechanisms.

## Acknowledgements

We thank Simon Sprecher’s and Claude Desplan’s lab for providing the mutant flies. We also like to thank Juusola lab members for fruitful discussions and comments.

## Author contributions

Conceptualisation, A.A.B., H.M., and M.J.; methodology, A.A.B., H.M. and M.J.; investigation, A.A.B. with H.M. performing ERG recordings; software, H.M. and J.K.; formal analysis, A.A.B. and H.M.; ERG recording analysis, M.J.; data curation, A.A.B.; visualisation, A.A.B..; writing – original draft, A.A.B., H.M.; writing – review and editing, A.A.B., H.M., J.K., and M.J.; resources, project administration, supervision, and funding acquisition, M.J.

## Methods

### Experimental model and study participant details

#### *Drosophila* husbandry and larval preparation

Experiments were performed using late third-instar *Drosophila melanogaster*. Approximately 80–100 adult flies were allowed to lay eggs for 12 h at 25°C on standard apple-juice agar plates supplemented with yeast paste. Plates were then maintained at 25°C for approximately 4 days, until the larvae reached the late third-instar stage. Before each experiment, larvae were collected from the culture plates, gently washed with distilled water to remove residual food, and transferred to Petri dishes containing 1% agarose. Behavioural experiments were conducted at room temperature in a darkened room to minimise uncontrolled visual stimulation. Each larva was used in only one experiment and was not reused across trials.

The fly lines used in behavioural and ERG experiments were the wild-type Canton-S line (WT) and the *Rh5*^*2*^ and *Rh6*^*1*^ mutants (provided by the Sprecher lab)e experiments and of the fly lines are shown in supplementary Table S1.

#### Experimental groups and replication

Behavioural assays were performed using independent groups of larvae. Each trial in the automated apparatus typically contained approximately 30 –50 larvae, and one larval group constituted one biological replicate. Naïve and trained larvae were tested in separate groups rather than repeatedly measuring the same animals. Statistical analyses were based on group-level behavioural choice counts, with experimental batch included as a random effect to account for shared experimental conditions. Sample sizes for each experimental condition are reported in the Results and Supplementary Table S1.

## Method details

### Automated behavioural apparatus

Larval colour-learning experiments were performed using a custom-built automated platform that integrated controlled visual stimulation, aversive reinforcement, motorised stimulus switching, and overhead behavioural recording within a single programmable system (Figure 1). The apparatus combined three-dimensionally printed structural components, commercially available electronic components, and custom control software. The printed components maintained reproducible alignment of the behavioural arena, illumination path, filter holders, rotating platform, and shock-delivery elements.

The behavioural arena consisted of a 90-mm Petri dish mounted above a motorised rotating platform. A broadband white-light source (See colour spectrum in **Figure S1**) positioned below the arena provided illumination through a diffusive plastic layer, which improved spatial uniformity across the agarose substrate. Interchangeable blue and green optical filters were positioned in the illumination path beneath the dish. Filters were selected from their measured transmission spectra to generate spectrally distinct blue and green conditions (See **Figure S1**).

An Arduino-controlled stepper motor rotated the filter platform to switch between colour conditions. An electronic relay synchronised platform movement and delivery of the aversive electrical stimulus. Light-source distance, illumination intensity, shock timing, stimulus duration, inter-epoch timing, and the number of training repetitions could be specified through the control software or adjusted mechanically.

### Spectral and intensity calibration

For the intensity-control experiments, reduced-intensity blue and green stimuli were generated by stacking two filters of the same spectral class used in the principal experiments. This manipulation reduced the light intensity to approximately one-third of that produced by the corresponding single-filter stimulus. The spectral profiles and relative intensities of the stimuli were measured using a CCS200/M compact spectrometer (Thorlabs).

### Initial colour-preference assay

Naïve larvae were placed near the centre of a fresh 90-mm Petri dish containing 1% agarose. The arena was illuminated from below so that one half was blue and the other half green (**Figure 1a**). Larvae moved freely for 60 s without electrical reinforcement. At the end of the assay, the numbers of larvae on the blue and green halves were counted and used to quantify baseline colour preference.

### Aversive colour-conditioning protocol

Larvae were trained under alternating blue and green illumination. One colour was designated as the reinforced conditioned stimulus (CS+) and paired with electrical shock, whereas the other served as the unreinforced conditioned stimulus (CS−). Blue and green were assigned reciprocally as CS+ across experimental groups. Each stimulus epoch lasted 30 s. Larvae underwent five CS+/CS− cycles, corresponding to ten 30-s training epochs in total.

The relay system controlled electrical reinforcement, and the Arduino controller synchronised it with CS+ presentation. The nominal controller setting was 110 V for 30s. Training was performed in a darkened room to minimise uncontrolled visual input. Electric shock was delivered via two copper wires embedded in the opposite side of the agarose in the Petri dish.

### Post-training colour-choice test

Immediately after training, larvae were transferred to a fresh Petri dish containing 1% agarose and divided into equal blue- and green-illuminated halves, matching the geometry of the initial-preference assay (**Figure 1b**). No electrical reinforcement was delivered. After 60 s, we counted the number of larvae on the two halves and used these counts to quantify post-training choice and calculate the behavioural index (see below). The orientation of the blue and green halves was randomised between trials by rotating the test arena relative to the apparatus and light source, thereby reducing fixed spatial and external-cue biases. Larvae and agarose plates were discarded after each experiment.

### Intensity-choice control assay

Although the blue and green filters were selected to produce closely matched light intensities, we performed an additional control experiment to determine whether aversive learning was driven by colour identity or by differences in stimulus intensity. Independent groups of larvae were trained using reduced-intensity blue or green stimuli generated by stacking filters of the same spectral class, producing about one-third of the light intensity generated by the corresponding single-filter stimulus.

During training, one colour was presented at approximately one-third of its original intensity, while the other remained at its original intensity. Blue and green were assigned reciprocally as the shock-associated conditioned stimulus (CS+). Following training, independent groups of larvae were tested under two intensity conditions: (i) an equalised-intensity condition, in which both colours were presented at their original intensities, and (ii) a reversed-intensity condition, in which the reduced-intensity cue was transferred to the alternative colour. This design allowed us to examine whether learned avoidance followed the spectral identity of the conditioned stimulus rather than the relative intensity relationship experienced during training.

If larvae learned the spectral identity of the shock-associated stimulus, they were expected to avoid the CS+ colour despite changes in relative stimulus intensity. Conversely, if avoidance depended solely on the reduced-intensity cue, preference was expected to follow that cue when transferred to the alternative colour. The training schedule and 60-s post-training choice test were otherwise identical to those used in the main experiment. Behaviour was quantified using the same conditioned-stimulus-aligned preference index described below.

### Extracellular recordings of larval visual responses

We recorded light-evoked extracellular field potentials from intact third-instar larvae using a preparation adapted from insect visual electrophysiology^42^. Although the larval visual organs are the Bolwig organs rather than compound eyes, the recording approach is methodologically comparable to that used for classical adult electroretinograms (ERGs). We therefore refer to these recordings as larval ERGs.

Larvae were anaesthetised with CO_2_ and immobilised on a glass slide using a minimal amount of low-melting wax, leaving the anterior head region exposed for recording (Figure 3b). This approach minimised movement during electrode placement while preserving the overall integrity of the animal. The use of wax-based immobilisation was based on standard adult *Drosophila* ERG procedures, in which rosin–beeswax mixtures or low-temperature melting wax are used to stabilise the preparation during ERG recording.

Recordings were made using pulled borosilicate glass electrodes filled with insect Ringer’s solution. Low-resistance sharp electrodes (approximately 1-10 MΩ) served as fine extracellular probes. A reference electrode was inserted into the posterior body wall, while the recording electrode was positioned over the anterior head region near the Bolwig organ/optic nerve entry zone under visual guidance with a micromanipulator (**Figure 3b**). We adjusted the recording electrode position until we obtained a stable light-evoked response.

UV, blue, green, amber and white (broadband) light stimuli were delivered using high-intensity LEDs, with the preparation housed inside a darkened, electrically shielded Faraday cage. The LED emission spectra are shown in **Figure 3b**. For each larva and stimulus condition, 10–20 repeated responses were recorded. Responses were sampled at 1 kHz and analysed offline in MATLAB. Repeated responses were baseline-corrected and averaged as described below to improve the signal-to-noise ratio.

### Electroretinogram signal processing and feature extraction

We analysed electroretinogram (ERG) recordings using custom-written scripts (Biosyst, Juusola^43-45^) in MATLAB (MathWorks, Natick, MA, USA). Individual responses were baseline-corrected by subtracting the mean voltage of the first 100 data points, corresponding to the 100-ms period before light stimulus onset. Corrected traces were then averaged to generate a mean response waveform for each larva and stimulus condition (**Figures 3c, d**).

We tested whether the temporal profile of the wild-type (Canton-S) ERG response could be reproduced by summing the Rh5^2^ and Rh6^1^ mutant ERG responses. Because recording amplitudes varied substantially with electrode position and larval muscle activity, we restricted this analysis to normalised waveform shapes rather than absolute response amplitudes.

For each LED stimulus, we calculated a mean response waveform across larvae for each genotype. Averaging reduced the contribution of sporadic muscle-activity-related noise. We then summed the mean Rh5^2^ and Rh6^1^ mutant waveforms and independently normalised the wild-type mean and the summed mutant waveform to a minimum of zero and a maximum of one. Residual waveforms were calculated by subtracting the normalised summed mutant waveform from the normalised wild-type waveform (**Figure 3d**).

### Quantification and statistical analysis

#### Behavioural indices

For associative assays, N_CS_ denotes the number of larvae choosing the colour paired with shock, N_other_ the number choosing the alternative colour, and N_total_ = N_CS_ + N_other_. For descriptive visualisation, the conditioned-colour preference index was:

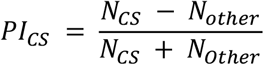

Positive PI_CS_ values indicate attraction of the shock-associated colour, values near zero indicate no conditioned bias, and negative values indicate relative avoidance of the conditioned colour. Naïve preference was visualised as:

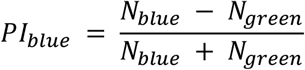

Positive PI_blue_ values indicate blue preference. These bounded indices were used for descriptive plots, not as the primary inferential responses.

#### Experimental unit and grouped-count inference

Counts from larvae tested together were analysed as one grouped two-choice observation. The total number of responding larvae contributed to the precision of that group’s estimate but did not increase the number of biological replicates. Innate and trained measurements came from independent groups and were not analysed as paired observations. Experimental batch was included as a blocking random effect because groups within a batch shared rearing and testing conditions.

#### Beta-binomial mixed-effects models

Primary inference used beta-binomial generalised linear mixed-effects models with a logit link. The beta-binomial distribution accommodates extra-binomial heterogeneity arising from shared group behaviour, arena variability, and residual differences among trials. For group i:

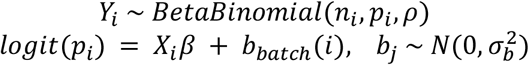

Parameters were estimated by maximum likelihood with a custom random-intercept likelihood integrated using 25-point Gauss–Hermite quadrature. Optimisation used multiple starting values and constrained maximum likelihood. Optimiser exit status, finite estimates, and the observed Hessian were checked. Grouped-binomial GLMMs were retained as sensitivity analyses, but beta-binomial models were used for the reported primary inference.

#### Standard blue-green experiment

The complete fixed-effect structure was training × genotype × conditioned colour, with batch as a random intercept. Training distinguished shock-conditioned from innate-control groups; genotype comprised wild type, Rh5, and Rh6; and conditioned colour distinguished blue from green CS assignments. Wild type, innate control, and blue CS were reference levels. Nested likelihood-ratio tests compared the complete model with models that removed the three-way and then all conditioned-colour interactions. Because the conditioned-colour interactions did not improve fit, planned contrasts were estimated from training × genotype + conditioned colour. Primary contrasts compared trained with innate groups within each genotype. Secondary difference-in-differences contrasts tested whether the training effect in each mutant was attenuated relative to wild type.

#### Innate blue-green preference

The response was N_blue_ out of N_blue_ + N_green_, with genotype as a fixed effect and batch as a random intercept. Genotype-specific marginal probabilities were tested against 0.5. Mutant-versus-wild-type contrasts were treated as control analyses because genotypes were tested in partly distinct experimental periods.

#### Wild-type colour-intensity experiment

The fixed factors were test mode (equalised or reversed), conditioned colour, and the colour presented at reduced intensity during training, with batch as a random intercept. A complete three-way model was compared with an additive model. Principal marginal tests estimated p(CS) in each test mode, averaging equally over conditioned colour and reduced-intensity assignment. The primary comparison tested reversed versus equalised choice.

#### Contrasts, multiplicity, and sensitivity analyses

Nested models were compared by likelihood-ratio tests. Planned contrasts used two-sided Wald z tests and are reported as ORs or marginal probabilities with 95% Wald CIs. Holm correction was applied separately within prespecified families: three within-genotype learning contrasts, two mutant-attenuation contrasts, three genotype-specific innate-preference tests, two innate genotype contrasts, and two intensity probability tests. The reversed-versus-equalised contrast was a single prespecified comparison. Statistical significance was defined as adjusted p < 0.05 within contrast families and p < 0.05 for single prespecified and model-selection tests. A sensitivity analysis replaced batch with experiment date as the random intercept.

### Significance threshold

Unless otherwise stated, statistical tests were two-sided and statistical significance was defined as p < 0.05.

### Lead contact

Further information and requests for resources should be directed to and will be fulfilled by the lead contact.

## Materials availability

This study did not generate new unique reagents.

## Data and code availability

Custom code used for apparatus control, behavioural analysis, statistical analysis, and three-dimensional design will be available at https://github.com/JuusolaLab/XXX after publication. Group-level source data underlying all main and supplemental figures should be deposited in the same repository or an archival repository and linked here before submission.

