## Supplemental data for "Associative learning reconfigures colour-guided behaviour in *Drosophila* larvae"

**SUPPLEMENTARY DATA**

Supplemental information includes five tables.

The numerical values below reproduce the final beta-binomial analysis supplied with the manuscript. Raw source data and executable code should accompany the submission repository.

**Table S1. Experimental design and sample sizes**

| **Experiment** | **Genotype** | **Groups** | **Batches** | **Responding larvae** | **Primary question** |
| --- | --- | --- | --- | --- | --- |
| Standard blue-green | Wild type | 40 | 35 across experiment | included in 3,971 | Training effect and colour adjustment |
| Standard blue-green | Rh5 | 32 | 35 across experiment | included in 3,971 | Within-genotype learning and attenuation vs WT |
| Standard blue-green | Rh6 | 32 | 35 across experiment | included in 3,971 | Within-genotype learning and attenuation vs WT |
| Colour-intensity control | Wild type | 60 | 15 | 2,171 | Does avoidance persist after intensity reversal? |

**Table S2. Planned contrasts from the standard beta-binomial model**

| **Contrast** | **OR** | **95% CI** | **Adjusted p** | **Interpretation** |
| --- | --- | --- | --- | --- |
| WT trained vs innate | 0.586 | 0.465–0.740 | 2.05 × 10⁻⁵ | Robust learned avoidance |
| Rh5 trained vs innate | 0.878 | 0.671–1.149 | 0.688 | No significant within-genotype effect |
| Rh6 trained vs innate | 1.008 | 0.776–1.310 | 0.950 | No detectable within-genotype effect |
| Rh5 attenuation vs WT | 1.497 | 1.049–2.137 | 0.026 | Training effect weaker than WT |
| Rh6 attenuation vs WT | 1.719 | 1.211–2.440 | 0.0048 | Training effect weaker than WT |
| Green vs blue CS adjustment | 1.194 | 1.029–1.386 | 0.019 | Modest colour asymmetry |

**Table S3. Genotype-specific innate blue choice**

| **Genotype** | **p(blue)** | **95% CI** | **Holm-adjusted conclusion** |
| --- | --- | --- | --- |
| Wild type | 0.488 | 0.449–0.527 | Not different from 0.5 |
| Rh5 | 0.460 | 0.411–0.511 | Not different from 0.5 |
| Rh6 | 0.508 | 0.460–0.555 | Not different from 0.5 |

**Table S4. Wild-type colour-intensity experiment**

| **Test/term** | **Estimate** | **95% CI** | **p value** | **Conclusion** |
| --- | --- | --- | --- | --- |
| Equalised test: p(CS) | 0.432 | 0.398–0.468 | 0.00042, Holm | Conditioned-colour choice below chance |
| Reversed test: p(CS) | 0.436 | 0.401–0.471 | 0.00042, Holm | Avoidance persists after cue transfer. |
| Reversed vs equalised OR | 1.014 | 0.829–1.239 | 0.896 | No effect of intensity reversal |
| Green vs blue CS OR | 1.283 | 1.049–1.569 | 0.031, Holm | Modest colour asymmetry |
| Reduced-intensity training colour OR | 1.048 | 0.855–1.285 | 0.652 | No detectable assignment effect |

**Table S5. Sensitivity to temporal clustering**

| **Contrast** | **Batch random intercept OR** | **Date random intercept OR** |
| --- | --- | --- |
| WT trained vs innate | 0.586 | 0.586 |
| Rh5 trained vs innate | 0.878 | 0.878 |
| Rh6 trained vs innate | 1.008 | 1.007 |
| Rh5 attenuation vs WT | 1.497 | 1.497 |
| Rh6 attenuation vs WT | 1.719 | 1.718 |

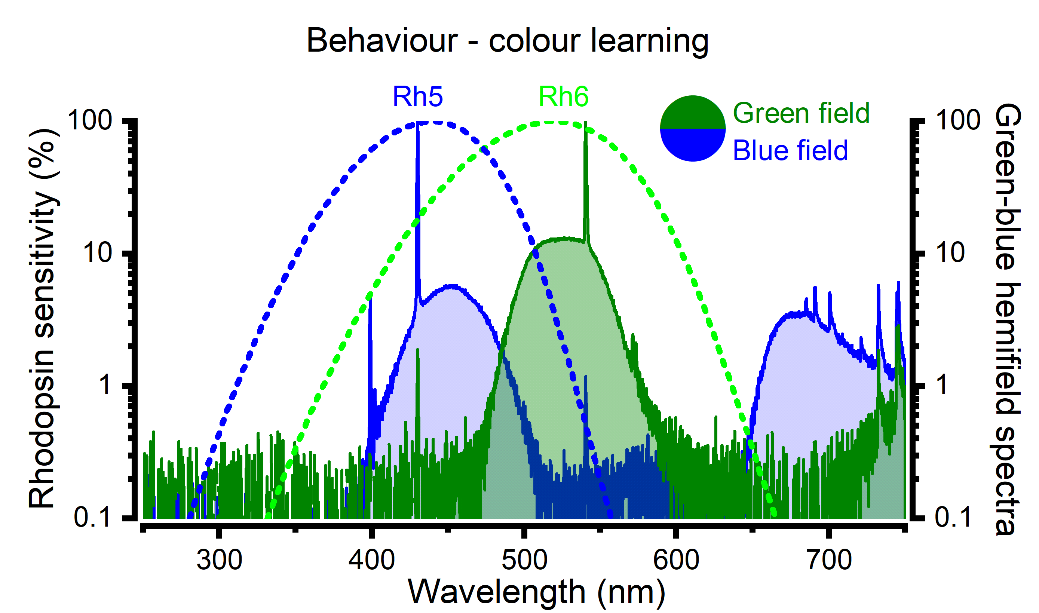

**Figure S1** The blue and green half-field stimulus spectra plotted with respect to the corresponding spectral sensitivity of larval Rh5 and Rh6 photoreceptors (nomograms).
